## Supplementary materials for "The Genetic Architecture of Multimodal Human Brain Age"

^*^Corresponding author:

**eMethod 1:** **The definition of genomic loci, independent significant SNP, lead SNP, candidate SNP**

**eMethod 2: Image quality check for MUSE**

**eText1: Exemplary genomic locus linked to GM, WM, and FC-BAG**

**eFigure 1:** **The five-layer neural network used for age prediction and its performance using WM-IDP**

**eFigure 2: Genetic correlation (*g_c_*) between the GM, WM, and FC-BAG using the LDSC software in the split-sample analyses**

**eFigure 3:** **Split-sample genome-wide association results**

**eFigure 4: Sex-stratified genome-wide association results**

**eFigure 5: Non-European genome-wide association results**

**eFigure 6:** **fastGWA for mixed linear models**

**eFigure 7: Machine learning-specific GWAS**

**eFigure 8: Feature type-specific GWAS**

**eFigure 9: Incremental R2 of the PRS derived by the PLINK C+T approach**

**eFigure 10:** **Results for the inverse Mendelian randomization for the seven clinical traits**

**eFigure 11: Sensitivity check for all other significant exposure variables in the forward MR analyses for A) cancer on GM-BAG, B) diabetes on GM-BAG, and C) AD on WM-BAG**

**eFigure 12:** **RNA expression overview of the *DNAJC1*** **gene in various cancer types**

**eTable 1: Brain age prediction performance using GM, WM, and FC-IDP**

**eTable 2: Identified genomic loci and mapped genes**

**eTable 3: Selected clinical traits for genetic correlations analyses**

**eTable 4: Results for genetic correlation estimates for the 16 clinical traits**

**eTable 5: Results for partitioned heritability estimates for the 53 functional categories (A) and cell type-specific analysis (B)**

**eTable 6: Selected exposure variables for the forward Mendelian randomization**

**eMethod 1:** **The definition of genomic loci, independent significant SNP, lead SNP, candidate SNP**

FUMA defined the significant independent SNPs, lead SNPs, candidate SNPs, and genomic risk loci as follows ([https://fuma.ctglab.nl/tutorial#snp2gene](https://fuma.ctglab.nl/tutorial%23snp2gene)):

*Independent significant SNPs*

They are defined as SNPs with *P*≤5×10^-8^ that are independent of each other at the user-defined *r^2^* (set to 0.6 in the current study). We further describe *candidate SNPs* as those in linkage disequilibrium (LD) with independent significant SNPs. FUMA then queries each candidate SNP in the GWAS Catalog to check whether any clinical traits have been reported to be associated with previous GWAS studies.

*Lead SNPs*

Lead SNPs are defined as independent significant SNPs that are also independent of each other at *r^2^*<0.1. If multiple independent significant SNPs are correlated at *r^2^*≥0.1, then the one with the lowest individual *P*-value becomes the lead SNP. If *r^2^* threshold is set to 0.1 for the independent significant SNPs, then they would constitute the identical set as the lead SNPs by definition. FUMA thus advises setting *r^2^* to be 0.6 or higher.

*Genomic risk loci*

FUMA defines genomic risk loci to include all independent signals physically close or overlapping in a single locus. First, independent significant SNPs dependent on each other at *r^2^*≥0.1 are assigned to the same genomic risk locus. Then, independent significant SNPs with less than the user-defined distance (250 kilobases by default) away from one another are merged into the same genomic risk locus - the distance between two LD blocks of two independent significant SNPs is the distance between the closest points from each LD block. Each locus is represented by the SNP within the locus with the lowest *P*-value.

**eMethod 2: Image quality check for MUSE**

T1-weighted MRIs were first quality checked (QC) for motion, image artifacts, or restricted field-of-view. Another QC was performed as follows: First, the images were examined by manually evaluating for pipeline failures (e.g., poor brain extraction, tissue segmentation, and registration errors). Furthermore, a second step automatically flagged images based on outlying values of quantified metrics; those flagged images were re-evaluated.

**eText1: Exemplary genomic locus linked to GM, WM, and FC-BAG**

The genomic locus (top lead SNP: rs534115641, **Fig. 2C**) linked to GM-BAG was mapped to multiple protein-encoding genes by position, eQTL, and chromatin interaction. The *NSF* gene, which encodes *N*-ethylmaleimide-sensitive fusion proteins, plays a key role in transferring membrane vesicles between cellular compartments. This gene has been linked to several conditions, including Parkinson's disease (PD)^1^, epithelial ovarian cancer^2^, cognitive traits^3^, and fibromuscular dysplasia^4^. The *CRHR1* gene encodes a G protein-coupled receptor, which specifically binds to neuropeptides of the corticotropin-releasing hormone family. These neuropeptides are recognized as key regulators of the hypothalamic-pituitary-adrenal pathway. A prior GWAS^5^ corroborated the association of this gene with the response to environmental stress, providing substantive support for the engagement of the hypothalamic-pituitary-adrenal axis, the central nervous system, and the endocrine system in the regulation of stress response^6^. We also identified a highly polygenic genomic locus (top lead SNP: rs564819152, **Fig. 2D**) for WM-BAG. This locus mapped to the *SKIDA1*, *CASC10*, *MLLT10*, and *DNAJC1* genes – all implicated in various types of cancer. In contrast, the FC-BAG locus was novel and did not map to any genes. All mapped genes for GM and WM-BAG are presented in **Supplementary eTable 2**.

**eFigure 1: The five-layer neural network used for age prediction and its performance using WM-IDP**

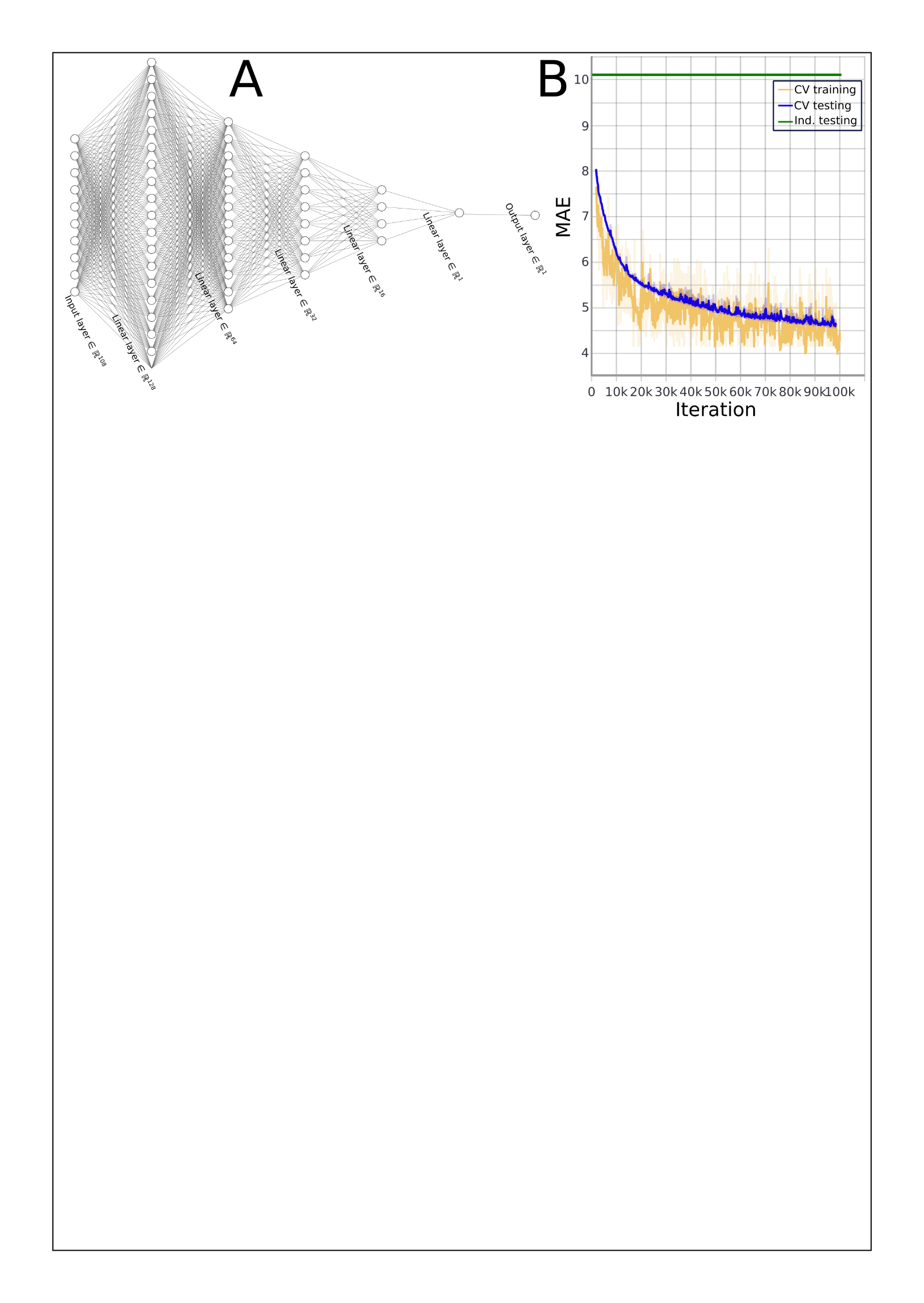

**A**) We illustrate the neural network architecture utilized in this study to predict brain age using GM, WM, and FC-IDP. The dimensionality of neurons in each linear layer is indicated in the diagram. **B**) We present the results of the cross-validation (CV) training, CV testing, and independent testing loss for the WM-IDP using FA, MD, ODI, and NDI from the TBSS-based approach. Notably, the network overfits the 108 WM-IDP since the number of network parameters (38,364) is significantly greater than the number of features (108). In addition, features from FA, MD, ODI, and NDI are highly correlated.

**eFigure 2: Genetic correlation (*g_c_*) between the GM, WM, and FC-BAG using the LDSC software in the split-sample analyses**

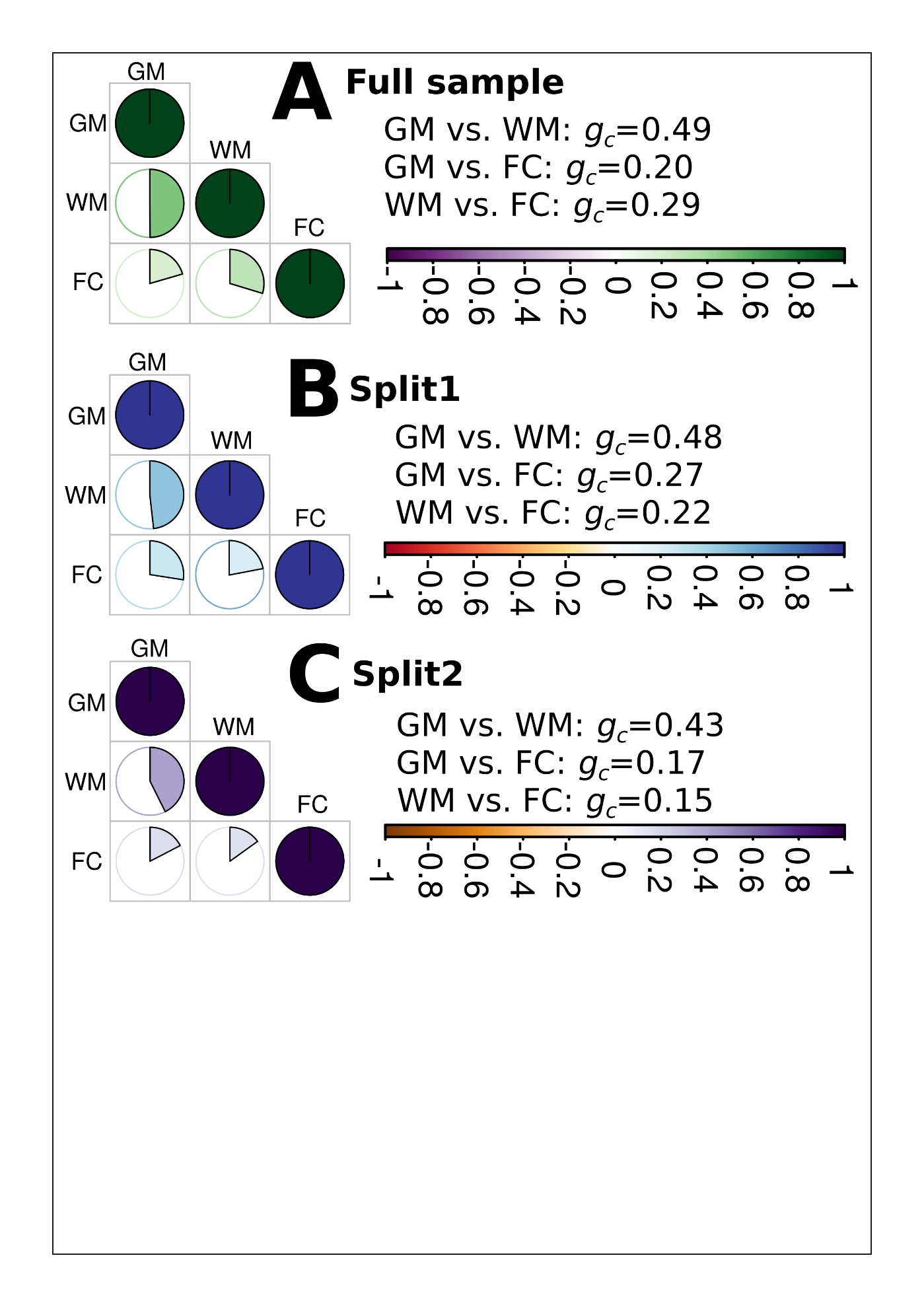

**A**) Genetic correlation using the full samples. **B**) Genetic correlation using the split1 sample. **C**) Genetic correlation using the split1 sample.

**eFigure 3: Split-sample genome-wide association results**

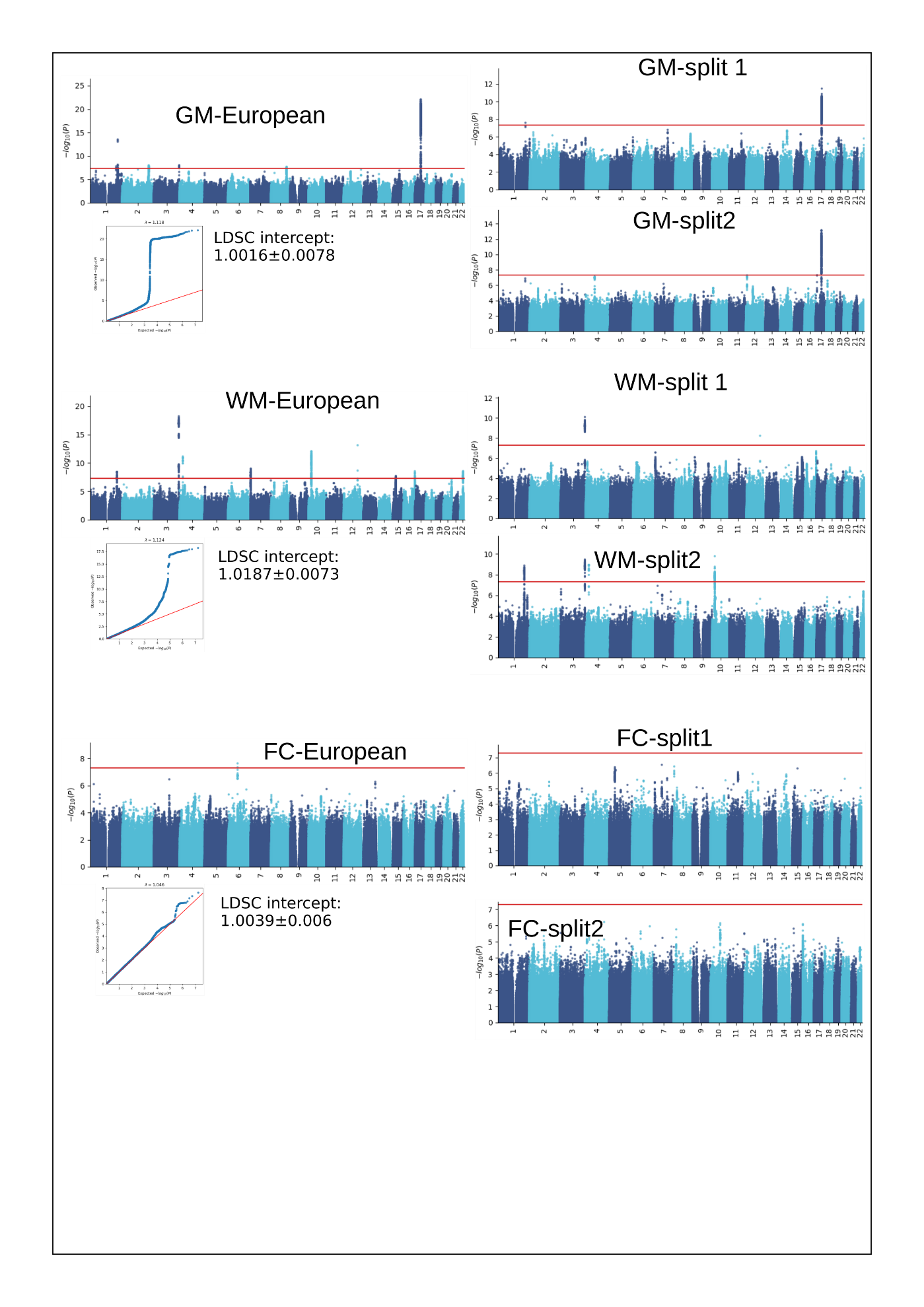

Genome-wide associations are presented for split-sample analyses (split1 vs. split2 vs. all). Genomic loci were identified using a genome-wide P-value threshold [–log_10_(P-value) > 7.30].

**eFigure 4: Sex-stratified genome-wide association results**

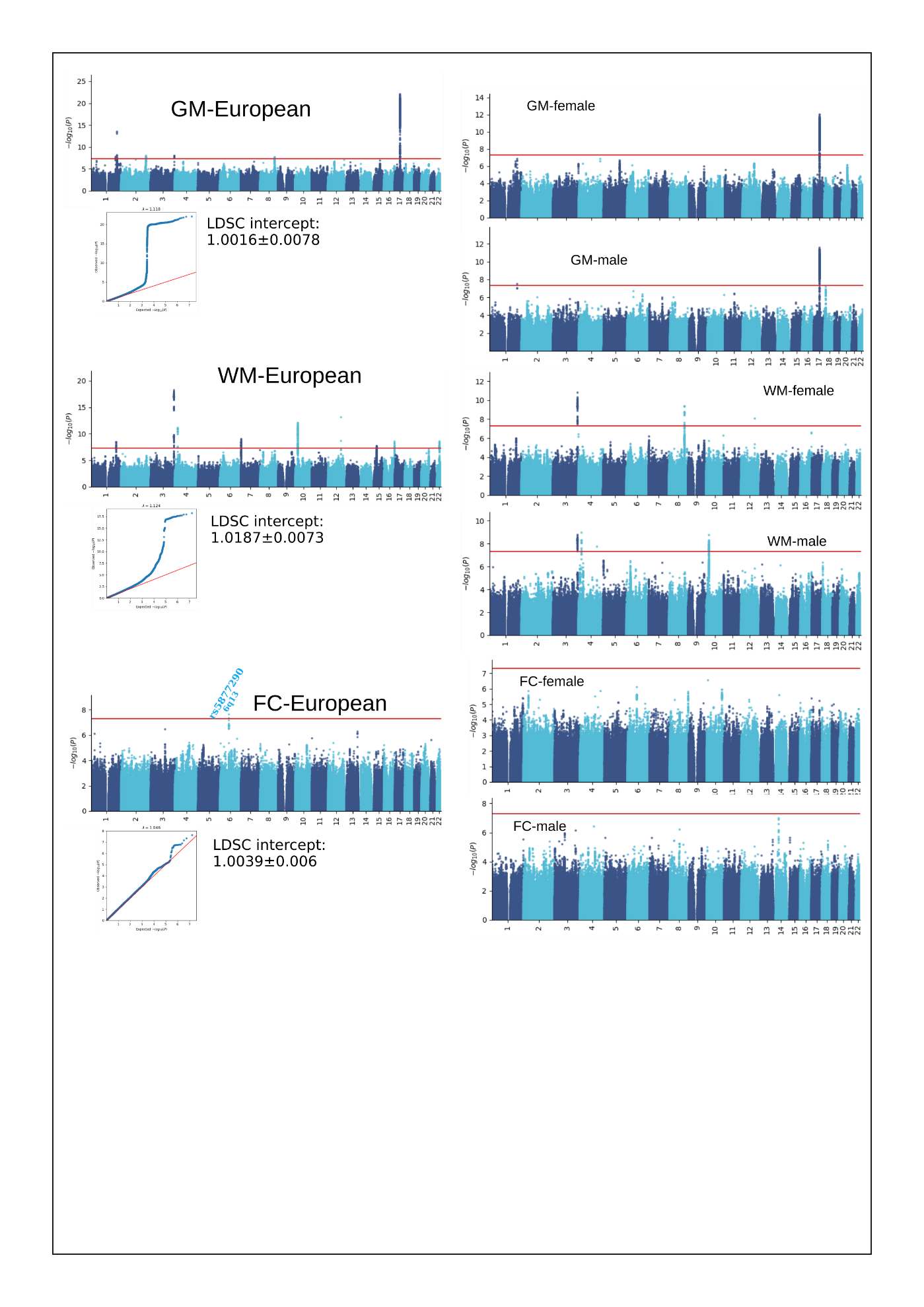

Genome-wide associations are presented for sex-stratified analyses (females vs. males). Genomic loci were identified using a genome-wide P-value threshold [–log_10_(P-value) > 7.30].

**eFigure 5: Non-European genome-wide association results**

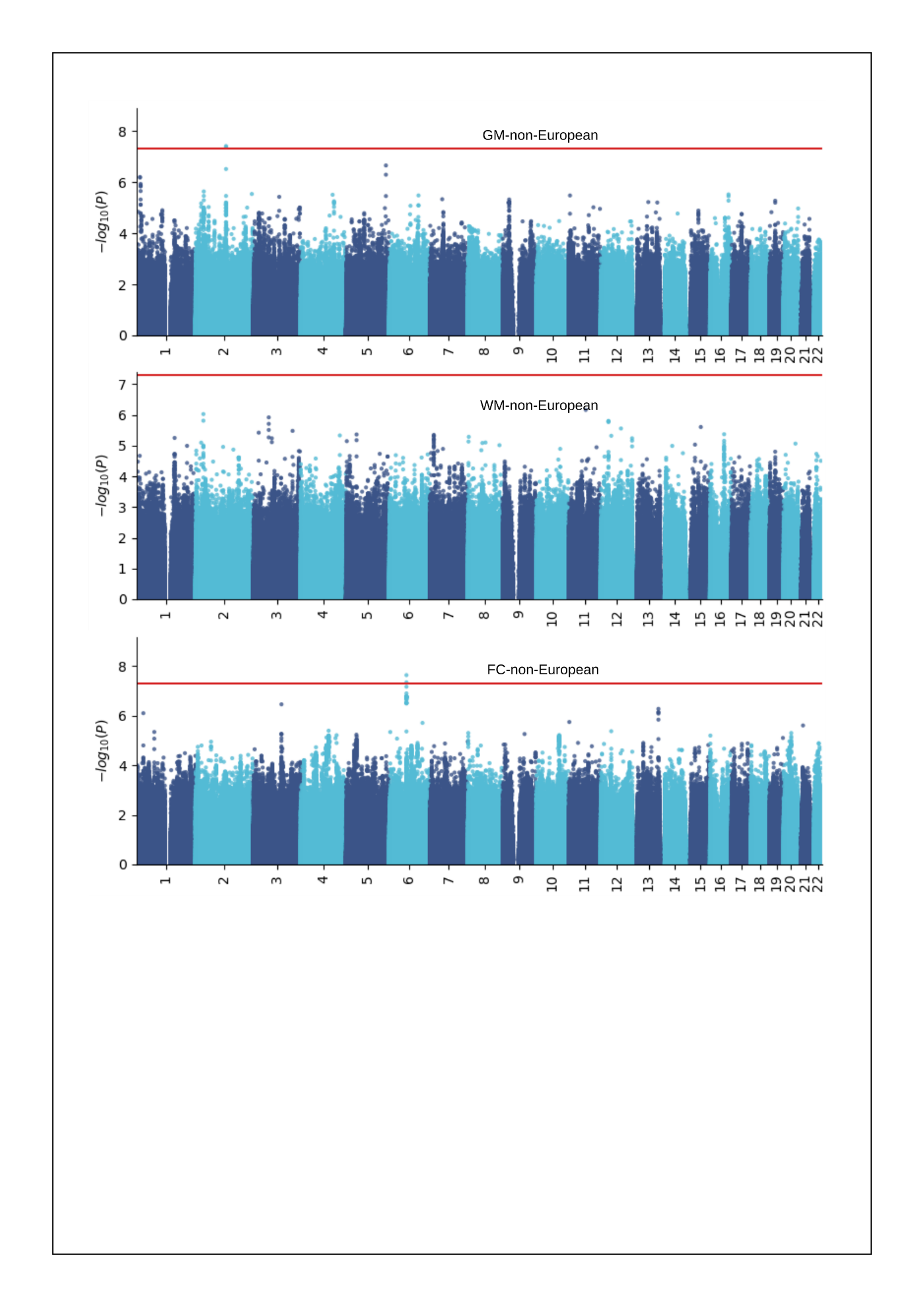

Genome-wide associations are presented for non-European populations in the UKBB study. Genomic loci associated were identified using a genome-wide P-value threshold [–log_10_(P-value) > 7.30].

**eFigure 6:** **fastGWA for mixed linear models**

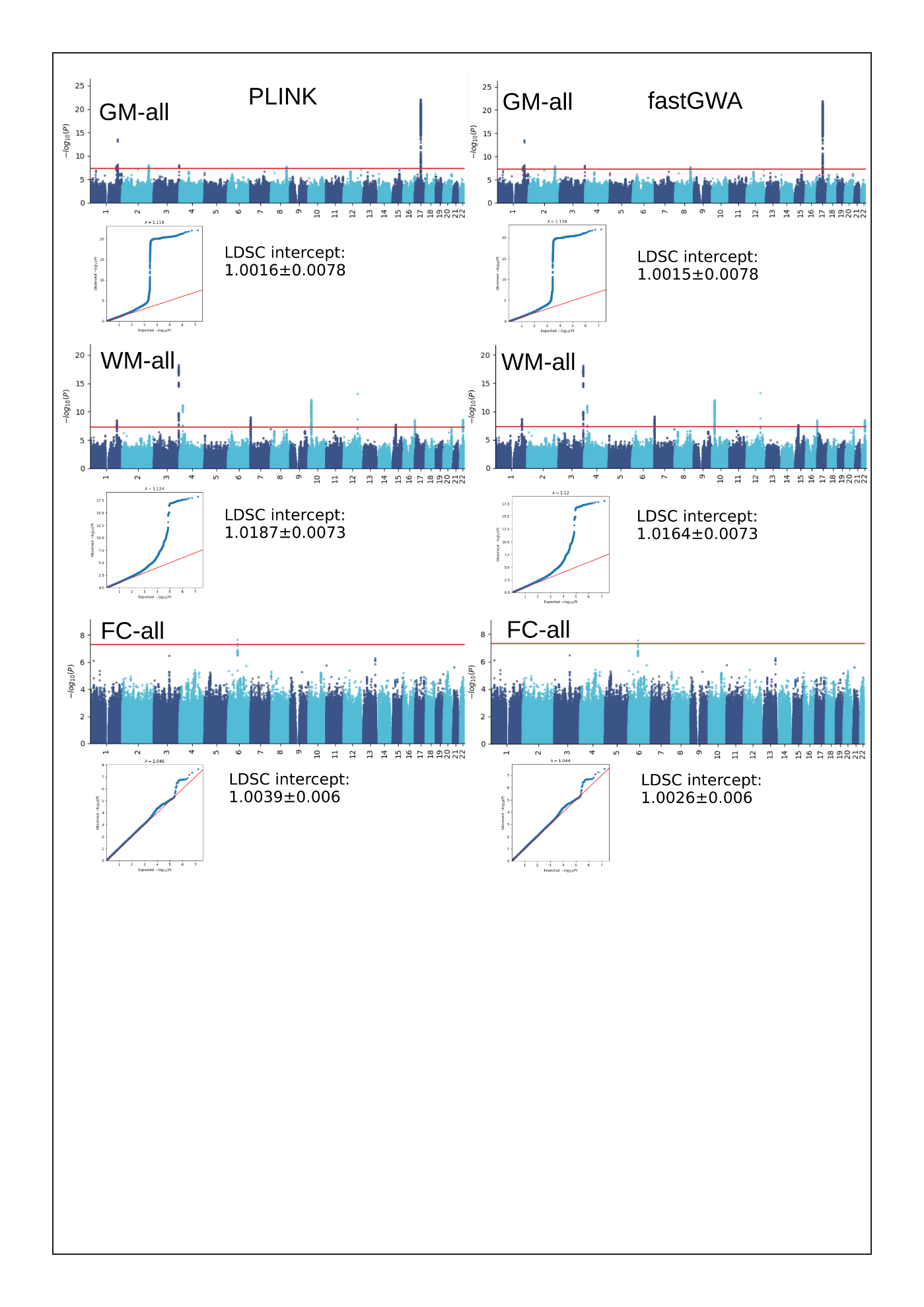

Genome-wide associations are presented for European populations in the UKBB study using fastGWA vs. PLINK. Genomic loci associated were identified using a genome-wide P-value threshold [–log_10_(P-value) > 7.30].

**eFigure 7: Machine learning-specific GWAS**

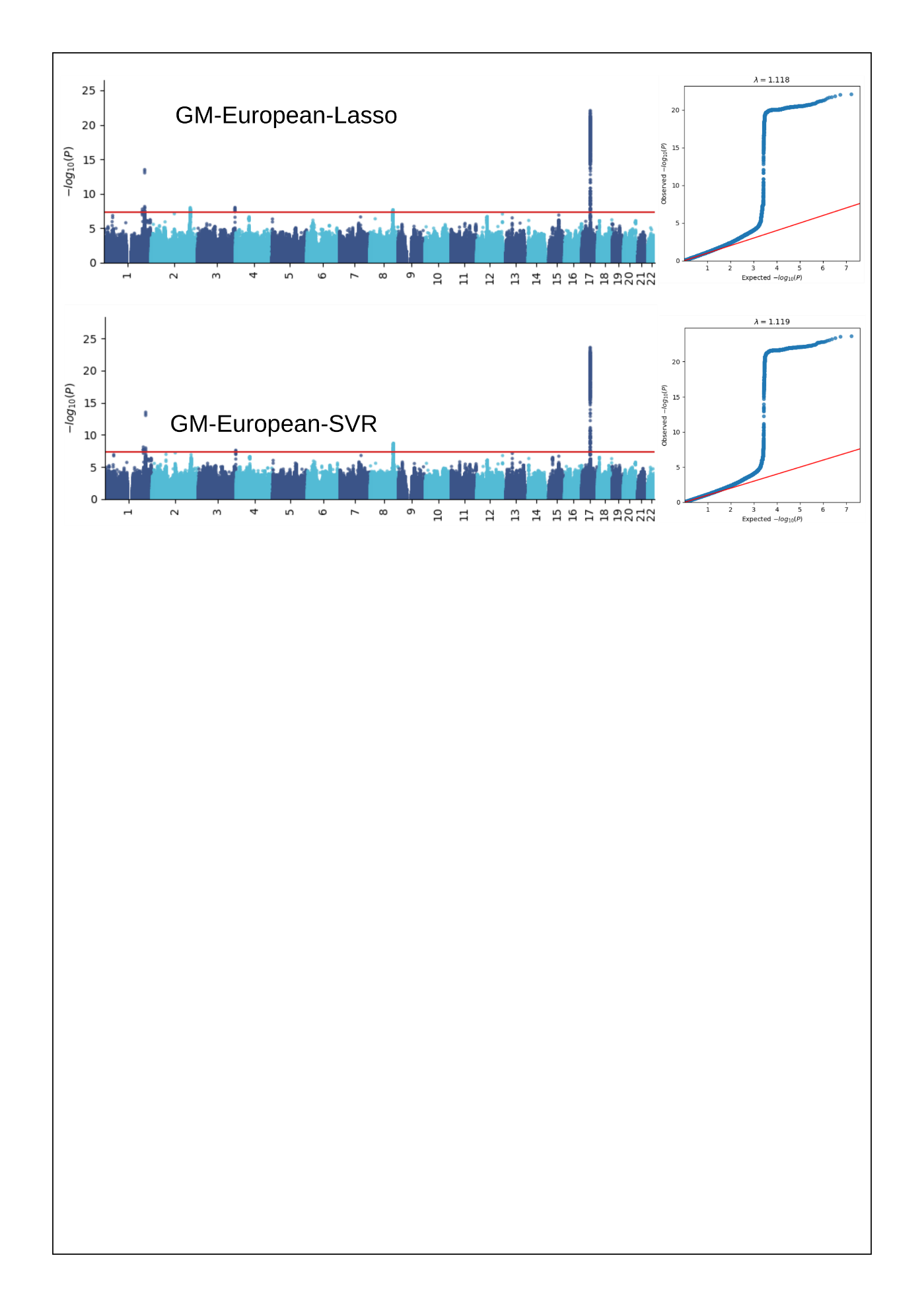

Genome-wide associations are presented for GM-BAG derived from Lasso regression (shown in the main text) and SVR. Genomic loci associated were identified using a genome-wide P-value threshold [–log_10_(P-value) > 7.30].

**eFigure 8: Feature type-specific GWAS**

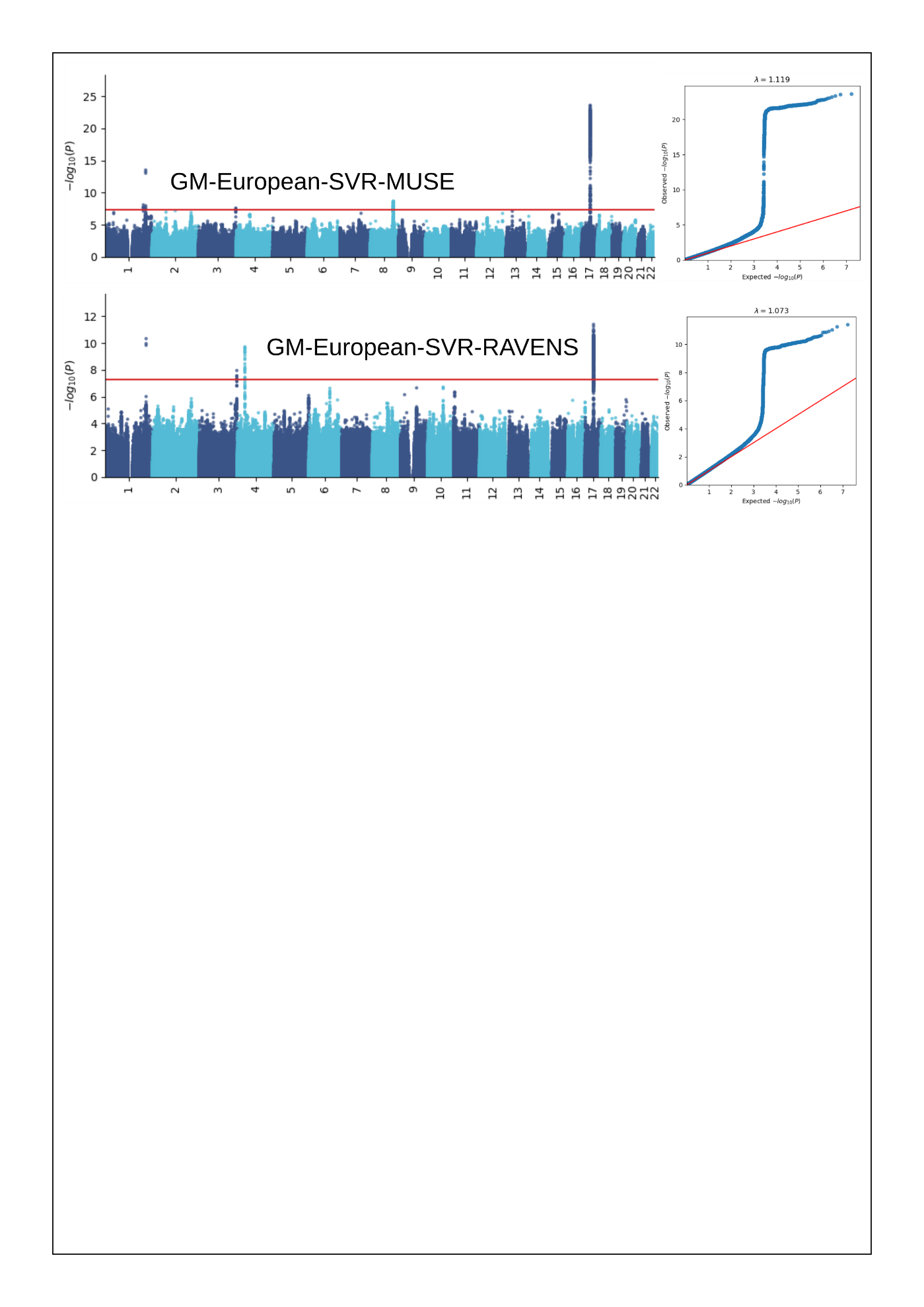

Genome-wide associations are presented for GM-BAG derived from MUSE ROIs (shown in the main text) and RAVENS voxel maps. Genomic loci associated were identified using a genome-wide P-value threshold [–log_10_(P-value) > 7.30].

**eFigure 9: Incremental R2 of the PRS derived by the PLINK C+T approach**

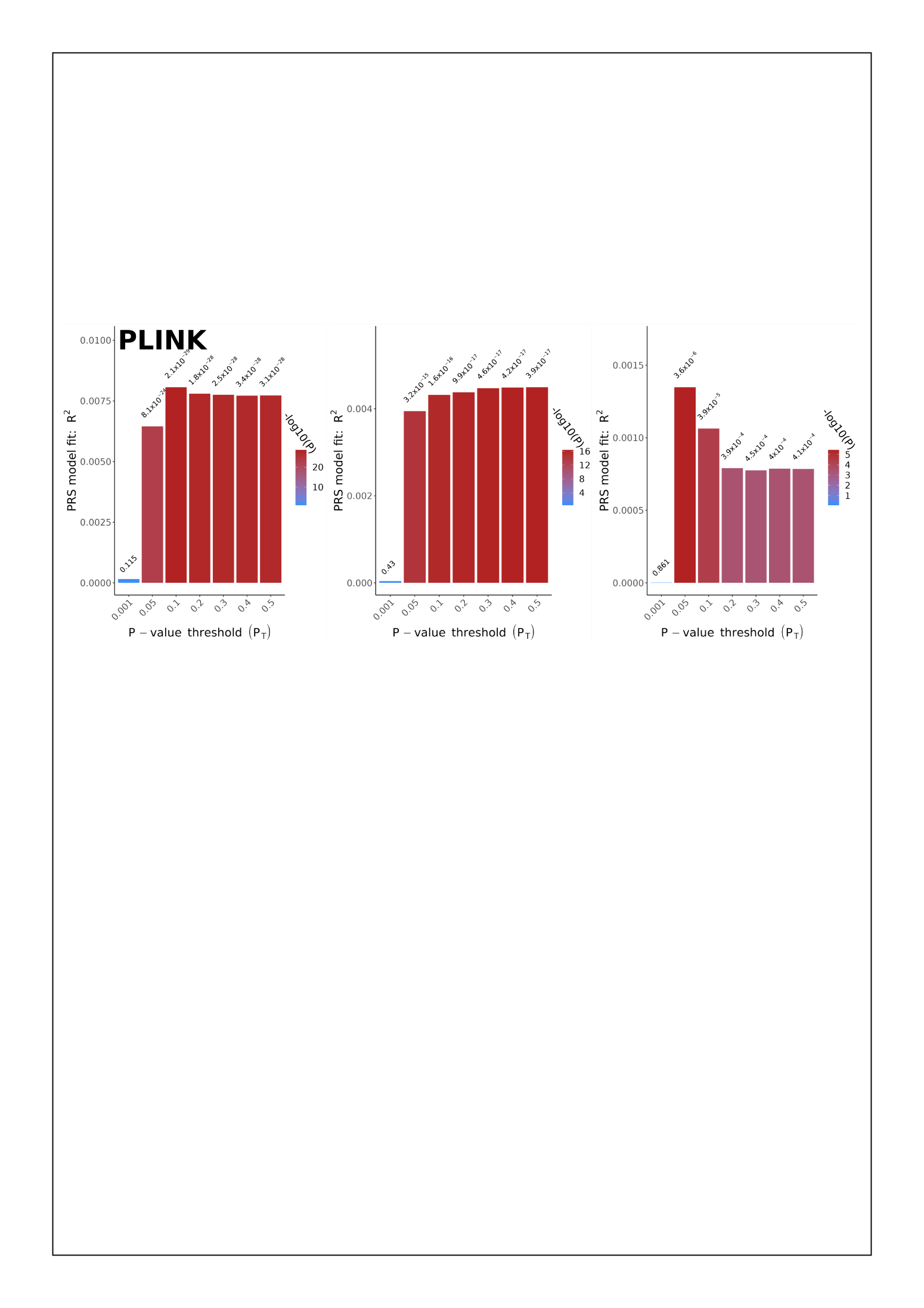

Incremental *R^2^* of the PRS derived by the PLINK C+T approach to predict the GM, WM, and FC-BAG in the target/test data (i.e., the split2 GWAS population in the split-sample analyses). The *y*-axis indicates the proportions of phenotypic variation (GM, WM, and FC-BAG) that the PRS can significantly and additionally explain. The *x*-axis lists the seven P-value thresholds considered.

**eFigure 10:** **Results for the inverse Mendelian randomization for the seven clinical traits**

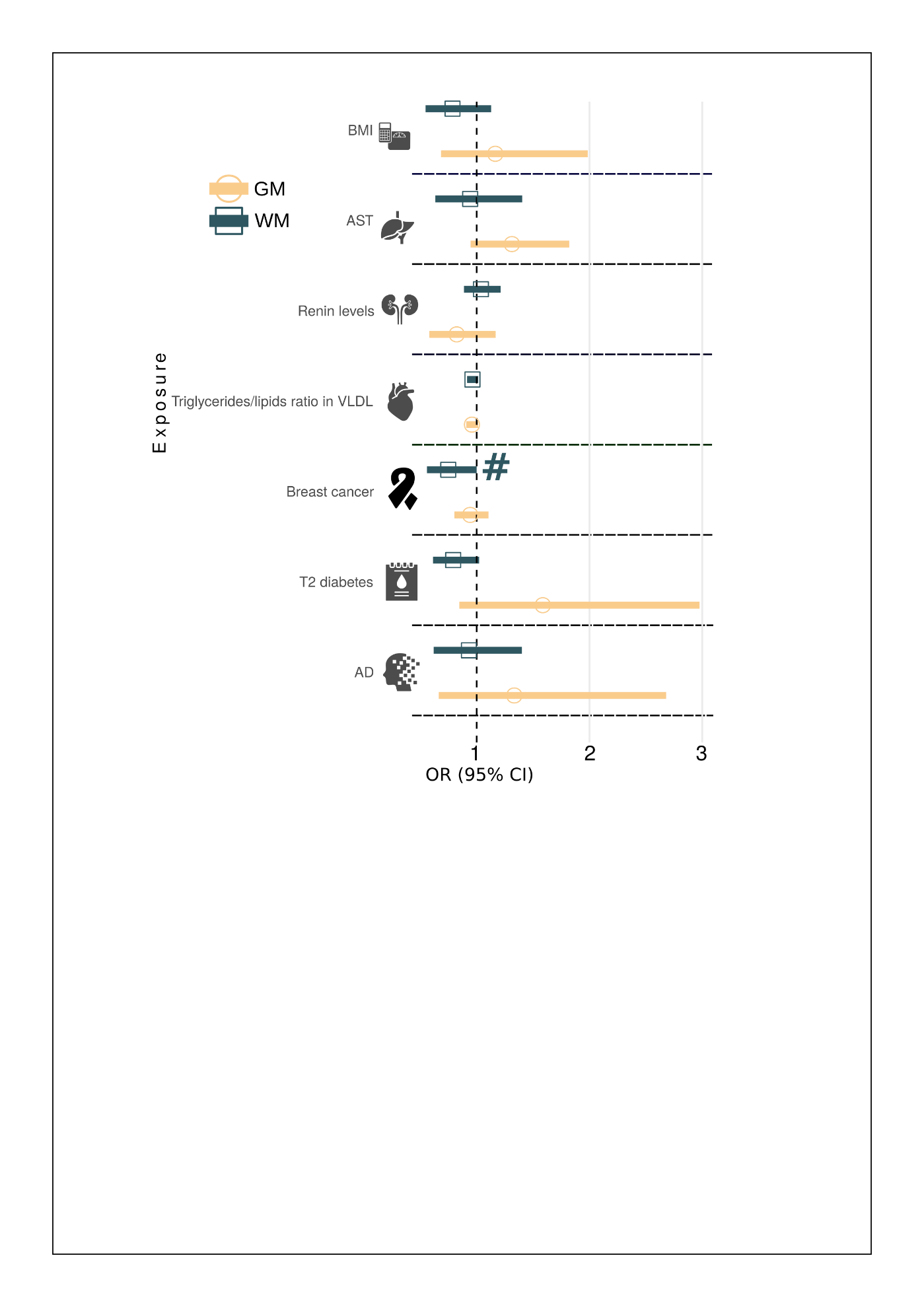

The inverse causal inference was performed using a two-sample Mendelian Randomization approach for seven selected clinical traits as outcome variables and GM, WM, and FC-BAG as exposure variables. Odds Ratio (OR) and its 95% confidence interval (CI) are presented. The symbol **#** indicates that the tests pass the nominal P-value threshold of 0.05 but do not survive the FDR correction. Abbreviation: AD: Alzheimer's Disease; AST: Aspartate Aminotransferase; BMI: Body Mass Index; VLDL: Very Low-Density Lipoprotein; SD: Standard Deviation; SE: Standard Error.

**eFigure 11: Sensitivity check for all other significant exposure variables in the forward MR analyses for 1) breast cancer on GM-BAG, 2) diabetes on GM-BAG, and 3) AD on WM-BAG.**

**1)** Sensitivity checks of causal effects of breast cancer on GM-BAG. **A**) Scatter plot indicates one potential outlier. **B**) Funnel plot shows no obvious asymmetry and points out two outliers. **C**) Single-SNP MR results. **D**) Leave-one-out analyses.

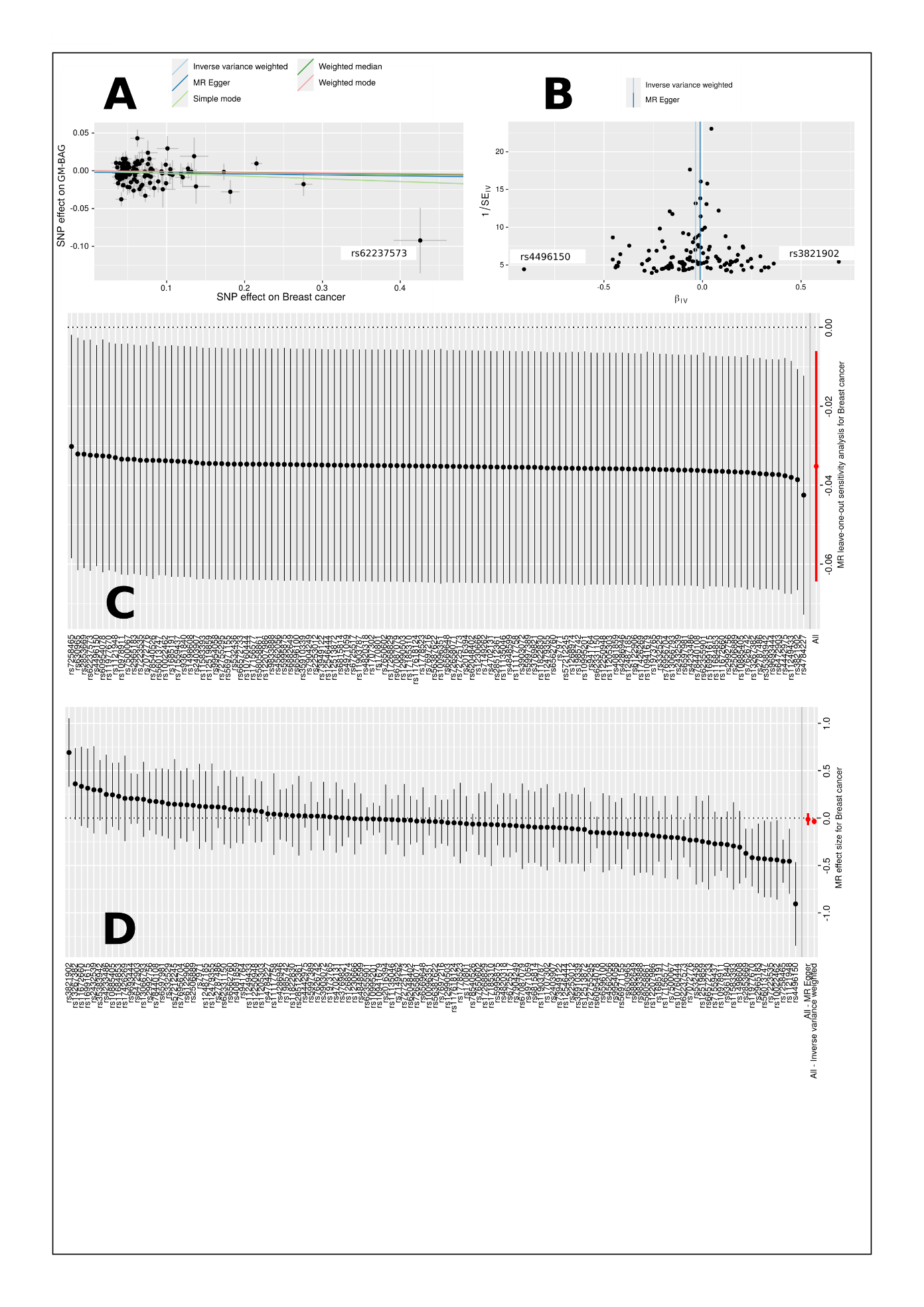

**2)** Sensitivity checks of causal effects of type 2 diabetes on GM-BAG. **A**) Scatter plot for the heterogeneity of the causal effects. **B**) Funnel plot shows the asymmetry of the causal effects. **C**) Single-SNP MR results. **D**) Leave-one-out analyses.

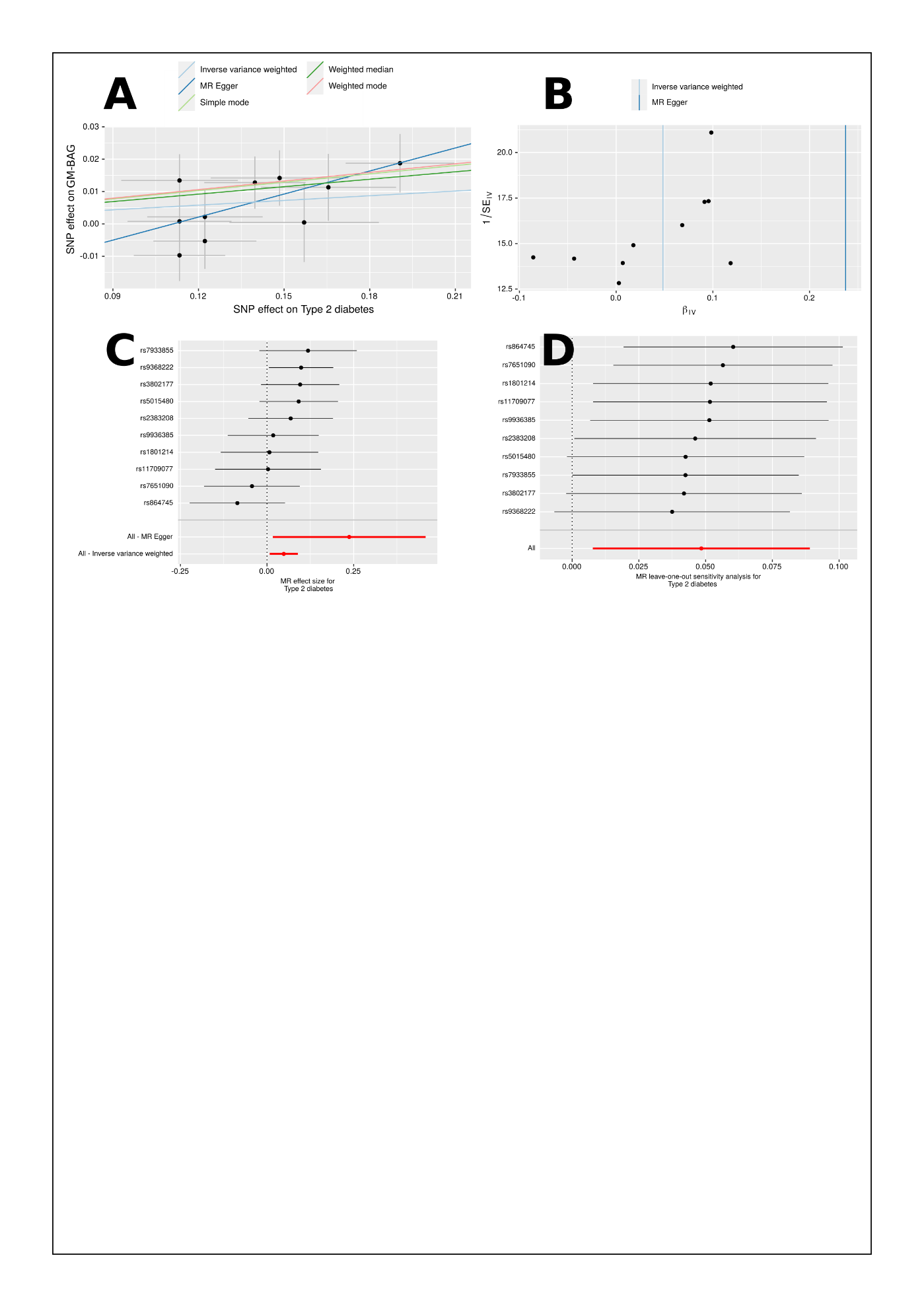

**3)** Sensitivity checks of causal effects of AD on WM-BAG. **A**) Scatter plot for the heterogeneity of the causal effects. **B**) Funnel plot shows no obvious asymmetry of the causal effects. **C**) Single-SNP MR results. **D**) Leave-one-out analyses.

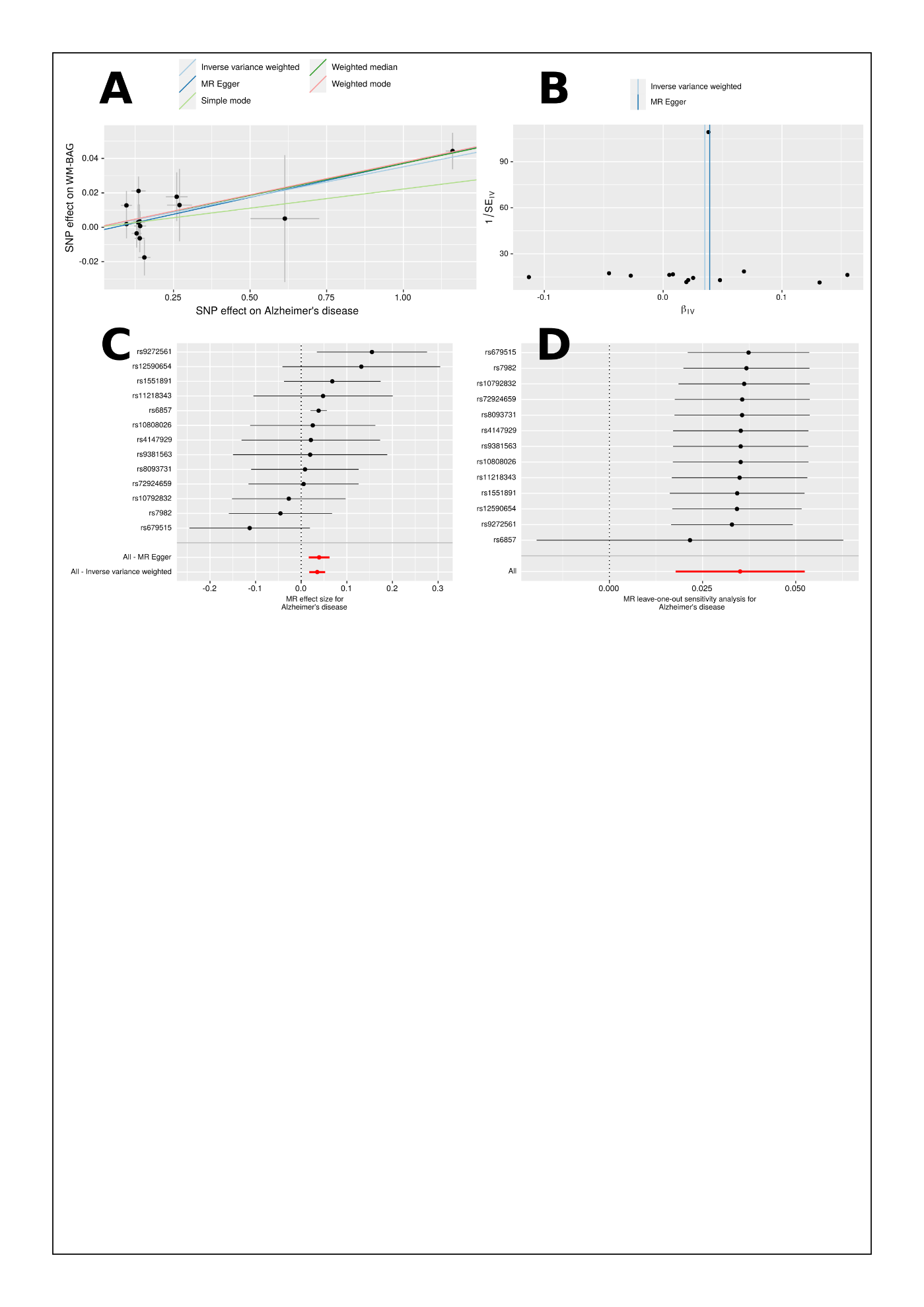

**eFigure 12:** **RNA expression overview of the *DNAJC1*** **gene in various cancer types.**

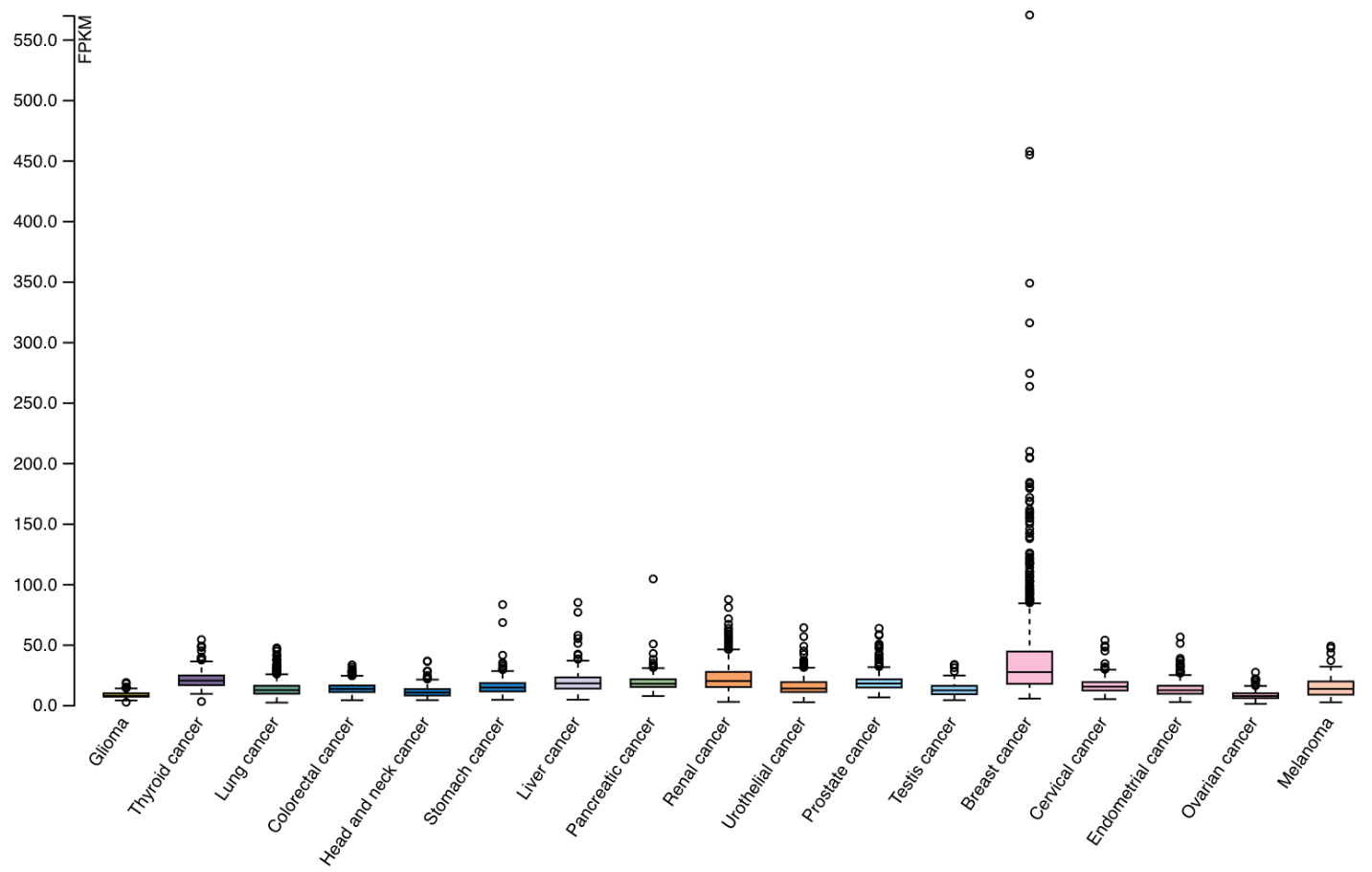

RNA expression overview shows RNA-seq data from The Cancer Genome Atlas (TCGA) project. FPKM represents fragments per kilobase of transcript per million mapped reads. The estimated gene expression values of the *DNAGC1* gene are displayed for 17 types of cancer/tumors.

**eTable 1: Brain age prediction performance using GM, WM, and FC-IDP.** We reported each machine learning model's mean absolute errors (MAE, year) and Pearson's correlation coefficient (r). The cross-validated (CV Test) and independent (Ind. Test) testing results were shown. Table **A** shows the results from the CV Test and the independent (Ind.) Test performance. Table **B** contains the results of the sex-stratified experiments.

**A**: **Brain age prediction results from the CV and independent test dataset**. The bolded text represents the lowest MAE for IDPs from each MRI modality. For WM-IDP, we fit the models with different combinations of features: i) 108 weighted mean TBSS WM-IDP from FA, MD, OD, and NDI; ii) 192 skeleton mean values of WM-IDP from FA, MD, OD, and NDI; iii) 48 FA WM-IDP.

| IDP | | Dataset | Linear SVR | | Lasso regression | | MLP | | NN | |
| --- | --- | --- | --- | --- | --- | --- | --- | --- | --- | --- |
|  |  |  | MAE | *r* | MAE | *r* | MAE | *r* | MAE | *r* |
| GM-IDP | | CV Test | 4.86 | 0.77 | 4.92 | 0.77 | 4.88 | 0.77 | 4.42 | 0.79 |
|  |  | Ind. Test | 4.43 | 0.66 | **4.39** | **0.66** | 5.35 | 0.64 | 4.83 | 0.65 |
| WM-IDP | 108 TBSS | CV Test | 4.88 | 0.77 | 4.94 | 0.78 | 5.29 | 0.75 | 4.54 | 0.79 |
|  |  | Ind. Test | 5.27 | 0.53 | 6.29 | 0.53 | 7.41 | 0.59 | 10.12 | 0.31 |
|  | 192 FA/MD/ODI/NDI | CV Test | 4.07 | 0.84 | 4.142 | 0.84 | 4.34 | 0.83 | 3.50 | 0.87 |
|  |  | Ind. Test | 21.90 | 0.71 | 21.66 | 0.73 | 6.12 | 0.71 | 15.77 | 0.30 |
|  | 48 FA | CV Test | 5.12 | 0.75 | 5.15 | 0.75 | 5.32 | 0.73 | 6.85 | 0.56 |
|  |  | Ind. Test | 5.02 | 0.65 | **4.92** | **0.65** | 7.95 | 0.60 | 6.84 | 0.42 |
| FC-IDP | | CV Test | 6.28 | 0.58 | 6.51 | 0.59 | 6.07 | 0.66 | 5.88 | 0.63 |
|  |  | Ind. Test | 5.97 | 0.43 | **5.48** | **0.44** | 6.02 | 0.46 | 6.05 | 0.43 |

**B**: **Brain age prediction results from the sex-stratified experiments.**

| IDP | Gender | Set | Linear SVR | | Lasso regression | | MLP | | NN | |
| --- | --- | --- | --- | --- | --- | --- | --- | --- | --- | --- |
|  |  |  | MAE | *r* | MAE | *r* | MAE | *r* | MAE | *r* |
| GM-IDP | Female | CV Test | 4.77 | 0.77 | 4.89 | 0.77 | 4.93 | 0.76 | 4.25 | 0.80 |
|  |  | Ind. Test | 4.46 | 0.64 | 4.44 | 0.64 | 6.94 | 0.60 | 5.02 | 0.62 |
|  | Male | CV Test | 4.69 | 0.78 | 4.77 | 0.79 | 4.70 | 0.79 | 4.08 | 0.82 |
|  |  | Ind. Test | 4.58 | 0.65 | 4.49 | 0.66 | 5.03 | 0.65 | 4.85 | 0.64 |
| WM-IDP | Female | CV Test | 4.81 | 0.78 | 4.77 | 0.79 | 4.73 | 0.78 | 5.61 | 0.66 |
|  |  | Ind. Test | 25.97 | 0.61 | 26.39 | 0.62 | 31.68 | 0.56 | 19.21 | 0.55 |
|  | Male | CV Test | 4.83 | 0.77 | 4.86 | 0.79 | 5.24 | 0.75 | 5.80 | 0.68 |
|  |  | Ind. Test | 7.88 | 0.63 | 5.67 | 0.62 | 10.24 | 0.62 | 14.96 | 0.56 |
| FC-IDP | Female | CV Test | 5.85 | 0.62 | 6.01 | 0.64 | 6.41 | 0.61 | 5.07 | 0.69 |
|  |  | Ind. Test | 5.93 | 0.42 | 5.58 | 0.42 | 5.36 | 0.42 | 6.28 | 0.41 |
|  | Male | CV Test | 6.03 | 0.60 | 6.32 | 0.62 | 6.83 | 0.59 | 6.11 | 0.62 |
|  |  | Ind. Test | 6.01 | 0.40 | 5.64 | 0.41 | 6.79 | 0.42 | 5.78 | 0.41 |

**eTable 2: Identified genomic loci and mapped genes.**

**GM-BAG:**

| Locus | Top lead SNP | P-value | Chromosome | Mapped genes |
| --- | --- | --- | --- | --- |
| 1 | rs61732315 | 1.63x10^-8^ | 1 | *MYOG, PPFIA4, ADORA1* |
| 2 | rs1452628 | 3.04x10^-14^ | 1 | *KCNK2, KCTD3* |
| 3 | rs186399184 | 1.05x10^-8^ | 2 | *CYP20A1, CARF, FAM117B, WDR12, ABI2, NBEAL1, ICA1L* |
| 4 | rs10933668 | 9.41x10^-9^ | 3 | NA |
| 5 | rs34051980 | 2.02x10^-8^ | 8 | *TNFRSF11B, COLEC10* |
| 6 | rs534115641 | 8.44x10^-23^ | 17 | *NSF, WNT3, KANSL1, CRHR1, NMT1, ARHGAP27, LRRC37A, EFCAB13, C17orf104, FMNL1, SPPL2C, ARL17A, MAPT, PLEKHM1, ARL17B, LRRC37A2, STH* |

**WM-BAG:**

| Locus | Top lead SNP | P-value | Chromosome | Mapped genes |
| --- | --- | --- | --- | --- |
| 1 | rs11118475 | 3.69x10^-09^ | 1 | *CD46, CR1L* |
| 2 | rs61067594 | 6.01x10^-19^ | 3 | *GMNC* |
| 3 | rs967140 | 7.26 x10^-12^ | 4 | *PPARGC1A* |
| 4 | rs2533872 | 9.95 x10^-10^ | 7 | *GNA12, AMZ1* |
| 5 | rs564819152 | 9.39 x10^-13^ | 10 | *SPAG6, MLLT10, DNAJC1, COMMD3, BMI1, SKIDA1, CASC10, COMMD3-BMI1* |
| 6 | rs12146713 | 7.67 x10^-14^ | 12 | *NUAK1* |
| 7 | rs654276 | 1.96 x10^-08^ | 15 | *TP53BP1, WDR76, ELL3, TUBGCP4, MFAP1, SERF2, ZSCAN29, TGM7, CASC4, CATSPER2, MAP1A, PDIA3, PPIP5K1, ADAL, LCMT2, FRMD5, SERINC4, CKMT1A, CKMT1B, HYPK, STRC, RP11-296A16.1, AC018512.1* |
| 8 | rs4843550 | 2.84 x10^-09^ | 16 | *C16orf95* |
| 9 | rs1894525 | 2.71x10^-09^ | 22 | *NOL12, TRIOBP, GCAT, ANKRD54, EIF3L, MICALL1, PICK1, GALR3, H1F0* |

**FC-BAG:**

| Locus | Top lead SNP | P-value | Chromosome | Mapped genes |
| --- | --- | --- | --- | --- |
| 1 | rs5877290 | 2.31x10^-8^ | 6 | NA |

**eTable 3: Selected clinical traits for genetic correlations analyses**. We selected the candidate studies from the GWAS Catalog for specific traits, including neurodegenerative diseases, psychiatric disorders, education, and intelligence. The inclusion criteria are i) GWAS summary statistics are publicly available; ii) the study population is European ancestry in the majority; iii) the heritability estimates (*h^2^*) via LDSC are not spuriously low (*h^2^*>0.05). This resulted in six clinical traits. We present the clinical trait, the dataset used, the URL link, the Pubmed ID, and the sample size. Abbreviations: PGC: Psychiatric genomics consortium; ADHD: attention deficit hyperactivity disorder; ASD: autism spectrum disorder; MDD: major depressive disorder; OCD: obsessive-compulsive disorder; SCZ: schizophrenia; BPD: bipolar disorder; SSGAC: Social Science Genetic Association Consortium; UKBB: UK Biobank.

| Trait | Dataset | URL | PubMed ID | Sample size |
| --- | --- | --- | --- | --- |
| AD | Meta | http://ftp.ebi.ac.uk/pub/databases/gwas | 30820047 | 63,926 |
| AD subtypes | UKBB | https://www.cbica.upenn.edu/bridgeport | NA | 33,540 |
| ADHD | PGC | <https://figshare.com/articles/dataset/adhd2019/14671965> | 30478444 | 53,293 |
| ASD | PGC | <https://figshare.com/articles/dataset/asd2019/14671989> | 30804558 | 46,350 |
| ASD subtypes | UKBB | <https://www.cbica.upenn.edu/bridgeport> | 37017948 | 14,786 |
| BPD | PGC | <https://figshare.com/articles/dataset/bip2019/14671998> | 31043756 | 51,710 |
| MDD | Meta | <https://figshare.com/articles/dataset/mdd2013/14672082> | 22472876 | 18,759 |
| Education | SSGAC | <http://ftp.ebi.ac.uk/pub/databases/gwas> | 23722424 | 126,559 |
| Intelligence | CTGlab | <http://ftp.ebi.ac.uk/pub/databases/gwas> | 28530673 | 78,308 |
| SCZ | PGC | <https://figshare.com/articles/dataset/scz2013sweden/14672154> | 23974872 | 11,244 |
| SCZ subtypes | UKBB | <https://www.cbica.upenn.edu/bridgeport> | 32103250 | 14,786 |
| OCD | Meta | https://figshare.com/articles/dataset/ocd2018/14672103 | 28761083 | 9,725 |

**eTable 4: Results for genetic correlation estimates for the 16 clinical traits**. We reported the genetic correlation (*r_g_*) estimates, their standard errors, and the Z and P-values. The details of the selected traits are presented in **eTable 3**. We also presented the h2 estimates using the LDSC software. Abbreviation: BAG: brain age gap.

| **BAG (*h^2^*)** | **Trait (*h^2^*)** | ***r_g_* mean** | ***r_g_* std** | **Z** | **P-value** |
| --- | --- | --- | --- | --- | --- |
| GM (0.30±0.03) | ADHD (0.23±0.01) | 0.0478 | 0.049 | 0.9767 | 0.3287 |
|  | AD (0.05±0.01) | 0.1595 | 0.1043 | 1.5293 | 0.1262 |
|  | AD1 (0.31±0.03) | 0.4001 | 0.0392 | 10.1976 | 2.03E-24 |
|  | AD2 (0.30±0.03) | -0.078 | 0.0517 | -1.5083 | 0.1315 |
|  | ASD (0.20±0.02) | 0.0718 | 0.0524 | 1.3685 | 0.1712 |
|  | ASD1 (0.27±0.03) | 0.3124 | 0.0671 | 4.6577 | 3.2E-06 |
|  | ASD2  (0.41±0.04) | -0.0713 | 0.057 | -1.2504 | 0.2111 |
|  | ASD3 (0.32±0.03) | -0.2078 | 0.0592 | -3.5096 | 0.0004 |
|  | BIP (0.34±0.02) | 0.0314 | 0.0389 | 0.8073 | 0.4195 |
|  | MDD (0.17±0.03) | 0.0637 | 0.0803 | 0.7932 | 0.4276 |
|  | OCD (0.34±0.04) | -0.1762 | 0.0665 | -2.6502 | 0.008 |
|  | SCZ (0.53±0.03) | 0.0543 | 0.0474 | 1.1458 | 0.2519 |
|  | SCZ1 (0.24±0.04) | 0.4908 | 0.0767 | 6.4017 | 1.54E-10 |
|  | SCZ2 (0.34±0.04) | 0.0879 | 0.0616 | 1.4271 | 0.1535 |
|  | Intelligence (0.19±0.01) | -0.068 | 0.0442 | -1.5379 | 0.1241 |
|  | Education (0.09±0.006) | -0.0764 | 0.0482 | -1.5853 | 0.1129 |
| WM (0.23±0.03) | ADHD (0.23±0.01) | 0.0478 | 0.049 | 0.9767 | 0.3287 |
|  | AD (0.05±0.01) | 0.0164 | 0.1381 | 0.1185 | 0.9057 |
|  | AD1 (0.31±0.03) | 0.2601 | 0.0472 | 5.5102 | 3.58E-08 |
|  | AD2 (0.30±0.03) | -0.0259 | 0.0531 | -0.4866 | 0.6265 |
|  | ASD (0.20±0.02) | 0.088 | 0.061 | 1.4419 | 0.1493 |
|  | ASD1 (0.27±0.03) | 0.339 | 0.0651 | 5.2076 | 1.91E-07 |
|  | ASD2  (0.41±0.04) | -0.012 | 0.0586 | -0.2045 | 0.8379 |
|  | ASD3 (0.32±0.03) | -0.0686 | 0.0756 | -0.9065 | 0.3647 |
|  | BIP (0.34±0.02) | 0.0466 | 0.043 | 1.0854 | 0.2777 |
|  | MDD (0.17±0.03) | 0.0291 | 0.0797 | 0.3649 | 0.7152 |
|  | OCD (0.34±0.04) | -0.1062 | 0.0723 | -1.4682 | 0.1421 |
|  | SCZ (0.53±0.03) | 0.1134 | 0.0496 | 2.2867 | 0.0222 |
|  | SCZ1 (0.24±0.04) | 0.4752 | 0.0995 | 4.7755 | 1.79E-06 |
|  | SCZ2 (0.34±0.04) | 0.2159 | 0.0665 | 3.2448 | 0.0012 |
|  | Intelligence (0.19±0.01) | -0.0802 | 0.0481 | -1.6668 | 0.0956 |
|  | Education (0.09±0.006) | 0.0118 | 0.0449 | 0.2616 | 0.7936 |
| FC (0.08±0.02) | ADHD (0.23±0.01) | 0.0478 | 0.049 | 0.9767 | 0.3287 |
|  | AD (0.05±0.01) | 0.3453 | 0.2031 | 1.7002 | 0.0891 |
|  | AD1 (0.31±0.03) | 0.2062 | 0.0762 | 2.7069 | 0.0068 |
|  | AD2 (0.30±0.03) | -0.0476 | 0.0842 | -0.5652 | 0.572 |
|  | ASD (0.20±0.02) | 0.0248 | 0.0935 | 0.2646 | 0.7913 |
|  | ASD1 (0.27±0.03) | 0.0204 | 0.1174 | 0.1735 | 0.8622 |
|  | ASD2  (0.41±0.04) | -0.0253 | 0.0895 | -0.2823 | 0.7777 |
|  | ASD3 (0.32±0.03) | 0.1325 | 0.1178 | 1.1253 | 0.2605 |
|  | BIP (0.34±0.02) | 0.0455 | 0.0673 | 0.6754 | 0.4994 |
|  | MDD (0.17±0.03) | 0.2776 | 0.1277 | 2.1731 | 0.0298 |
|  | OCD (0.34±0.04) | 0.1232 | 0.1179 | 1.045 | 0.296 |
|  | SCZ (0.53±0.03) | 0.0914 | 0.0756 | 1.21 | 0.2263 |
|  | SCZ1 (0.24±0.04) | 0.5042 | 0.1667 | 3.0243 | 0.0025 |
|  | SCZ2 (0.34±0.04) | 0.025 | 0.1119 | 0.2233 | 0.8233 |
|  | Intelligence (0.19±0.01) | -0.0544 | 0.0699 | -0.7792 | 0.4359 |
|  | Education (0.09±0.006) | -0.2231 | 0.0915 | -2.4393 | 0.0147 |

**eTable 5: Results for partitioned heritability estimates for the 53 functional categories (A) and cell type-specific analysis (B).** We reported the partitioned heritability estimates, the standard errors, and the Z and P-values. In total, there are 52 categories presented here, as we did not show the results for all available SNPs (the baseline model) for Table A.

**A: 52 functional categories**

| **IDP** | | **Category** | | **Prop.SNPs** | **Prop.h2** | **Prop.h2 std error** | **Enrichment** | **Enrichment std error** | **Enrichment p** |
| --- | --- | --- | --- | --- | --- | --- | --- | --- | --- |
| GM | Coding_UCSC_0 | | | 0.014658 | 0.079259 | 0.0403 | 5.407178 | 2.749352 | 0.112994 |
|  | Coding_UCSC.extend.500_0 | | | 0.064555 | 0.142005 | 0.043945 | 2.199733 | 0.68074 | 0.081496 |
|  | Conserved_LindbladToh_0 | | | 0.026063 | 0.427339 | 0.068669 | 16.3967 | 2.634763 | 5.8E-09 |
|  | Conserved_LindbladToh.extend.500_0 | | | 0.332514 | 0.573709 | 0.087212 | 1.725366 | 0.262279 | 0.005605 |
|  | CTCF_Hoffman_0 | | | 0.023829 | 0.113062 | 0.064653 | 4.74476 | 2.71324 | 0.171835 |
|  | CTCF_Hoffman.extend.500_0 | | | 0.071062 | 0.180246 | 0.075716 | 2.53645 | 1.065487 | 0.153483 |
|  | DGF_ENCODE_0 | | | 0.137594 | 0.546304 | 0.163117 | 3.970416 | 1.185499 | 0.011573 |
|  | DGF_ENCODE.extend.500_0 | | | 0.541501 | 0.897579 | 0.100433 | 1.657576 | 0.185472 | 0.000347 |
|  | DHS_peaks_Trynka_0 | | | 0.111766 | 0.440296 | 0.138217 | 3.939456 | 1.236663 | 0.017239 |
|  | DHS_Trynka_0 | | | 0.167755 | 0.41416 | 0.160964 | 2.46884 | 0.95952 | 0.125129 |
|  | DHS_Trynka.extend.500_0 | | | 0.498779 | 0.666078 | 0.125032 | 1.335417 | 0.250677 | 0.183209 |
|  | Enhancer_Andersson_0 | | | 0.004335 | 0.016029 | 0.032195 | 3.697193 | 7.426098 | 0.716007 |
|  | Enhancer_Andersson.extend.500_0 | | | 0.019069 | 0.108518 | 0.036138 | 5.690784 | 1.895097 | 0.013374 |
|  | Enhancer_Hoffman_0 | | | 0.06332 | 0.176 | 0.088002 | 2.779517 | 1.389797 | 0.197096 |
|  | Enhancer_Hoffman.extend.500_0 | | | 0.153929 | 0.286682 | 0.082588 | 1.862428 | 0.53653 | 0.114418 |
|  | FetalDHS_Trynka_0 | | | 0.084757 | 0.234658 | 0.124082 | 2.768617 | 1.463986 | 0.229531 |
|  | FetalDHS_Trynka.extend.500_0 | | | 0.28501 | 0.424903 | 0.117569 | 1.490834 | 0.412507 | 0.231563 |
|  | H3K27ac_Hnisz_0 | | | 0.391169 | 0.751898 | 0.057897 | 1.922184 | 0.14801 | 3.45E-10 |
|  | H3K27ac_Hnisz.extend.500_0 | | | 0.422591 | 0.810579 | 0.05623 | 1.918119 | 0.13306 | 1.46E-10 |
|  | H3K27ac_PGC2_0 | | | 0.269477 | 0.555377 | 0.09844 | 2.060948 | 0.365302 | 0.004046 |
|  | H3K27ac_PGC2.extend.500_0 | | | 0.336034 | 0.723373 | 0.074661 | 2.152675 | 0.222182 | 1.14E-06 |
|  | H3K4me1_peaks_Trynka_0 | | | 0.171318 | 0.779147 | 0.143365 | 4.547955 | 0.836838 | 4.28E-05 |
|  | H3K4me1_Trynka_0 | | | 0.426568 | 1.069136 | 0.125945 | 2.506366 | 0.295252 | 1.37E-07 |
|  | H3K4me1_Trynka.extend.500_0 | | | 0.609157 | 0.936822 | 0.057847 | 1.5379 | 0.094962 | 2.8E-07 |
|  | H3K4me3_peaks_Trynka_0 | | | 0.041789 | 0.304871 | 0.092573 | 7.295418 | 2.21523 | 0.004324 |
|  | H3K4me3_Trynka_0 | | | 0.133307 | 0.397945 | 0.088425 | 2.985168 | 0.663314 | 0.002171 |
|  | H3K4me3_Trynka.extend.500_0 | | | 0.255482 | 0.491154 | 0.085863 | 1.922461 | 0.336082 | 0.006558 |
|  | H3K9ac_peaks_Trynka_0 | | | 0.03877 | 0.299865 | 0.095064 | 7.734434 | 2.451996 | 0.00702 |
|  | H3K9ac_Trynka_0 | | | 0.126111 | 0.547579 | 0.091954 | 4.342035 | 0.72915 | 7.19E-06 |
|  | H3K9ac_Trynka.extend.500_0 | | | 0.230583 | 0.603484 | 0.082295 | 2.617204 | 0.356897 | 2.64E-05 |
|  | Intron_UCSC_0 | | | 0.387453 | 0.494738 | 0.042022 | 1.276898 | 0.108456 | 0.009974 |
|  | Intron_UCSC.extend.500_0 | | | 0.39713 | 0.557698 | 0.034255 | 1.404321 | 0.086256 | 3.63E-06 |
|  | PromoterFlanking_Hoffman_0 | | | 0.008427 | 0.041987 | 0.041434 | 4.982225 | 4.916611 | 0.41439 |
|  | PromoterFlanking_Hoffman.extend.500_0 | | | 0.033471 | 0.086502 | 0.053758 | 2.584347 | 1.606076 | 0.322961 |
|  | Promoter_UCSC_0 | | | 0.031164 | 0.048568 | 0.040694 | 1.558486 | 1.305799 | 0.669112 |
|  | Promoter_UCSC.extend.500_0 | | | 0.038627 | 0.020869 | 0.031134 | 0.540274 | 0.806022 | 0.567672 |
|  | Repressed_Hoffman_0 | | | 0.461222 | 0.227695 | 0.110905 | 0.493677 | 0.24046 | 0.034595 |
|  | Repressed_Hoffman.extend.500_0 | | | 0.719053 | 0.394491 | 0.054223 | 0.548626 | 0.075409 | 2.09E-09 |
|  | SuperEnhancer_Hnisz_0 | | | 0.16842 | 0.368778 | 0.034387 | 2.189631 | 0.204175 | 6.41E-09 |
|  | SuperEnhancer_Hnisz.extend.500_0 | | | 0.171607 | 0.384836 | 0.035614 | 2.242545 | 0.207533 | 5.58E-09 |
|  | TFBS_ENCODE_0 | | | 0.132451 | 0.396207 | 0.139937 | 2.991355 | 1.056519 | 0.059315 |
|  | TFBS_ENCODE.extend.500_0 | | | 0.343444 | 0.636668 | 0.109095 | 1.853779 | 0.317651 | 0.008761 |
|  | Transcribed_Hoffman_0 | | | 0.345418 | 0.405626 | 0.103077 | 1.174303 | 0.298412 | 0.557053 |
|  | Transcribed_Hoffman.extend.500_0 | | | 0.763062 | 0.710194 | 0.063587 | 0.930717 | 0.083331 | 0.412599 |
|  | TSS_Hoffman_0 | | | 0.018219 | 0.15972 | 0.051569 | 8.766743 | 2.830526 | 0.008052 |
|  | TSS_Hoffman.extend.500_0 | | | 0.034826 | 0.141052 | 0.047034 | 4.050251 | 1.350556 | 0.027411 |
|  | UTR_3_UCSC_0 | | | 0.011054 | 0.076751 | 0.030058 | 6.943103 | 2.719097 | 0.028483 |
|  | UTR_3_UCSC.extend.500_0 | | | 0.026931 | 0.084798 | 0.032526 | 3.148689 | 1.207734 | 0.07742 |
|  | UTR_5_UCSC_0 | | | 0.005425 | 0.008818 | 0.022127 | 1.62567 | 4.079002 | 0.878187 |
|  | UTR_5_UCSC.extend.500_0 | | | 0.027806 | 0.059455 | 0.036045 | 2.138208 | 1.296312 | 0.385518 |
|  | WeakEnhancer_Hoffman_0 | | | 0.021093 | 0.054834 | 0.065996 | 2.599675 | 3.128883 | 0.608383 |
|  | WeakEnhancer_Hoffman.extend.500_0 | | | 0.088958 | 0.100055 | 0.065093 | 1.124738 | 0.731722 | 0.864864 |
| WM | | | Coding_UCSC_0 | 0.014658 | 0.155365 | 0.04994 | 10.59922 | 3.406953 | 0.005012 |
|  |  |  | Coding_UCSC.extend.500_0 | 0.064555 | 0.1789 | 0.057994 | 2.771257 | 0.898359 | 0.051425 |
|  |  |  | Conserved_LindbladToh_0 | 0.026063 | 0.322703 | 0.088628 | 12.38191 | 3.400594 | 0.001085 |
|  |  |  | Conserved_LindbladToh.extend.500_0 | 0.332514 | 0.787166 | 0.086135 | 2.367317 | 0.259042 | 5.18E-07 |
|  |  |  | CTCF_Hoffman_0 | 0.023829 | -0.07361 | 0.079892 | -3.08904 | 3.352728 | 0.215354 |
|  |  |  | CTCF_Hoffman.extend.500_0 | 0.071062 | 0.088353 | 0.095646 | 1.243318 | 1.345942 | 0.856634 |
|  |  |  | DGF_ENCODE_0 | 0.137594 | 0.463824 | 0.188755 | 3.370972 | 1.371831 | 0.085516 |
|  |  |  | DGF_ENCODE.extend.500_0 | 0.541501 | 0.998602 | 0.122523 | 1.844137 | 0.226265 | 0.000339 |
|  |  |  | DHS_peaks_Trynka_0 | 0.111766 | 0.510709 | 0.197265 | 4.569458 | 1.764986 | 0.043526 |
|  |  |  | DHS_Trynka_0 | 0.167755 | 0.44245 | 0.209805 | 2.637481 | 1.250664 | 0.189588 |
|  |  |  | DHS_Trynka.extend.500_0 | 0.498779 | 0.892095 | 0.124579 | 1.788559 | 0.249768 | 0.002934 |
|  |  |  | Enhancer_Andersson_0 | 0.004335 | 0.013879 | 0.043332 | 3.20128 | 9.99505 | 0.825393 |
|  |  |  | Enhancer_Andersson.extend.500_0 | 0.019069 | 0.056812 | 0.045398 | 2.979252 | 2.380722 | 0.403264 |
|  |  |  | Enhancer_Hoffman_0 | 0.06332 | 0.12957 | 0.112742 | 2.046268 | 1.780508 | 0.558749 |
|  |  |  | Enhancer_Hoffman.extend.500_0 | 0.153929 | 0.443049 | 0.094994 | 2.878267 | 0.617128 | 0.003456 |
|  |  |  | FetalDHS_Trynka_0 | 0.084757 | 0.29676 | 0.164787 | 3.50132 | 1.944238 | 0.198713 |
|  |  |  | FetalDHS_Trynka.extend.500_0 | 0.28501 | 0.751411 | 0.113738 | 2.636434 | 0.399065 | 0.000152 |
|  |  |  | H3K27ac_Hnisz_0 | 0.391169 | 0.789498 | 0.068352 | 2.018306 | 0.174738 | 3.12E-09 |
|  |  |  | H3K27ac_Hnisz.extend.500_0 | 0.422591 | 0.867771 | 0.073537 | 2.053453 | 0.174014 | 7.33E-09 |
|  |  |  | H3K27ac_PGC2_0 | 0.269477 | 0.683085 | 0.121377 | 2.534857 | 0.450416 | 0.000673 |
|  |  |  | H3K27ac_PGC2.extend.500_0 | 0.336034 | 0.699731 | 0.083563 | 2.082321 | 0.248673 | 3.39E-05 |
|  |  |  | H3K4me1_peaks_Trynka_0 | 0.171318 | 0.595128 | 0.179485 | 3.473819 | 1.047668 | 0.018314 |
|  |  |  | H3K4me1_Trynka_0 | 0.426568 | 0.69677 | 0.13662 | 1.633432 | 0.320276 | 0.049134 |
|  |  |  | H3K4me1_Trynka.extend.500_0 | 0.609157 | 0.931012 | 0.066936 | 1.528362 | 0.109884 | 1.08E-05 |
|  |  |  | H3K4me3_peaks_Trynka_0 | 0.041789 | 0.054389 | 0.104481 | 1.301493 | 2.50017 | 0.904041 |
|  |  |  | H3K4me3_Trynka_0 | 0.133307 | 0.486131 | 0.107918 | 3.646695 | 0.809546 | 0.001314 |
|  |  |  | H3K4me3_Trynka.extend.500_0 | 0.255482 | 0.592693 | 0.094446 | 2.319899 | 0.369678 | 0.000614 |
|  |  |  | H3K9ac_peaks_Trynka_0 | 0.03877 | 0.352943 | 0.101732 | 9.103484 | 2.623986 | 0.002697 |
|  |  |  | H3K9ac_Trynka_0 | 0.126111 | 0.520868 | 0.112355 | 4.130231 | 0.890923 | 0.000588 |
|  |  |  | H3K9ac_Trynka.extend.500_0 | 0.230583 | 0.740526 | 0.09121 | 3.211533 | 0.395564 | 4.69E-08 |
|  |  |  | Intron_UCSC_0 | 0.387453 | 0.413057 | 0.052748 | 1.066084 | 0.136141 | 0.625957 |
|  |  |  | Intron_UCSC.extend.500_0 | 0.39713 | 0.507811 | 0.038444 | 1.278702 | 0.096804 | 0.003134 |
|  |  |  | PromoterFlanking_Hoffman_0 | 0.008427 | -0.03312 | 0.053711 | -3.93042 | 6.373536 | 0.439526 |
|  |  |  | PromoterFlanking_Hoffman.extend.500_0 | 0.033471 | 0.096027 | 0.064002 | 2.868915 | 1.912139 | 0.330485 |
|  |  |  | Promoter_UCSC_0 | 0.031164 | 0.180979 | 0.059716 | 5.807366 | 1.916211 | 0.012537 |
|  |  |  | Promoter_UCSC.extend.500_0 | 0.038627 | 0.156695 | 0.045908 | 4.05661 | 1.1885 | 0.009738 |
|  |  |  | Repressed_Hoffman_0 | 0.461222 | 0.176248 | 0.128626 | 0.382132 | 0.27888 | 0.026509 |
|  |  |  | Repressed_Hoffman.extend.500_0 | 0.719053 | 0.432354 | 0.059519 | 0.601283 | 0.082774 | 1.79E-06 |
|  |  |  | SuperEnhancer_Hnisz_0 | 0.16842 | 0.423493 | 0.044166 | 2.514504 | 0.262237 | 6.05E-09 |
|  |  |  | SuperEnhancer_Hnisz.extend.500_0 | 0.171607 | 0.462015 | 0.044715 | 2.692291 | 0.260564 | 9.87E-10 |
|  |  |  | TFBS_ENCODE_0 | 0.132451 | 0.315911 | 0.169577 | 2.385124 | 1.280304 | 0.283593 |
|  |  |  | TFBS_ENCODE.extend.500_0 | 0.343444 | 0.673625 | 0.128517 | 1.961386 | 0.374202 | 0.014155 |
|  |  |  | Transcribed_Hoffman_0 | 0.345418 | 0.522131 | 0.123845 | 1.511591 | 0.358535 | 0.144861 |
|  |  |  | Transcribed_Hoffman.extend.500_0 | 0.763062 | 0.656892 | 0.081823 | 0.860864 | 0.10723 | 0.201496 |
|  |  |  | TSS_Hoffman_0 | 0.018219 | 0.149603 | 0.053922 | 8.211426 | 2.959671 | 0.016901 |
|  |  |  | TSS_Hoffman.extend.500_0 | 0.034826 | 0.147209 | 0.0598 | 4.227051 | 1.71713 | 0.063105 |
|  |  |  | UTR_3_UCSC_0 | 0.011054 | 0.118996 | 0.032486 | 10.76476 | 2.938823 | 0.000811 |
|  |  |  | UTR_3_UCSC.extend.500_0 | 0.026931 | 0.145899 | 0.036224 | 5.417473 | 1.345067 | 0.001271 |
|  |  |  | UTR_5_UCSC_0 | 0.005425 | 0.093648 | 0.02992 | 17.26384 | 5.515599 | 0.004242 |
|  |  |  | UTR_5_UCSC.extend.500_0 | 0.027806 | 0.078239 | 0.037726 | 2.813762 | 1.35675 | 0.185159 |
|  |  |  | WeakEnhancer_Hoffman_0 | 0.021093 | -0.0458 | 0.079391 | -2.17161 | 3.763938 | 0.397779 |
|  |  |  | WeakEnhancer_Hoffman.extend.500_0 | 0.088958 | 0.120023 | 0.080342 | 1.349207 | 0.903146 | 0.699612 |
| FC | | | Coding_UCSC_0 | 0.014658 | 0.141202 | 0.181769 | 9.632997 | 12.40057 | 0.485702 |
|  |  |  | Coding_UCSC.extend.500_0 | 0.064555 | 0.083034 | 0.198569 | 1.286238 | 3.075949 | 0.925664 |
|  |  |  | Conserved_LindbladToh_0 | 0.026063 | 0.437223 | 0.295482 | 16.77593 | 11.33743 | 0.156162 |
|  |  |  | Conserved_LindbladToh.extend.500_0 | 0.332514 | 0.91434 | 0.344908 | 2.74978 | 1.037275 | 0.071678 |
|  |  |  | CTCF_Hoffman_0 | 0.023829 | -0.02302 | 0.296973 | -0.966 | 12.46275 | 0.872976 |
|  |  |  | CTCF_Hoffman.extend.500_0 | 0.071062 | 0.20233 | 0.279516 | 2.847222 | 3.933397 | 0.639695 |
|  |  |  | DGF_ENCODE_0 | 0.137594 | -0.51347 | 0.703396 | -3.7318 | 5.112124 | 0.332741 |
|  |  |  | DGF_ENCODE.extend.500_0 | 0.541501 | 0.699476 | 0.456609 | 1.291736 | 0.843229 | 0.731578 |
|  |  |  | DHS_peaks_Trynka_0 | 0.111766 | -0.1752 | 0.635589 | -1.56755 | 5.68679 | 0.64777 |
|  |  |  | DHS_Trynka_0 | 0.167755 | 0.530218 | 0.670341 | 3.160668 | 3.995956 | 0.588556 |
|  |  |  | DHS_Trynka.extend.500_0 | 0.498779 | 0.208197 | 0.489707 | 0.417413 | 0.981813 | 0.534689 |
|  |  |  | Enhancer_Andersson_0 | 0.004335 | -0.08131 | 0.131213 | -18.7558 | 30.26566 | 0.506936 |
|  |  |  | Enhancer_Andersson.extend.500_0 | 0.019069 | -0.06524 | 0.167418 | -3.42149 | 8.779532 | 0.605279 |
|  |  |  | Enhancer_Hoffman_0 | 0.06332 | 0.180646 | 0.375325 | 2.852895 | 5.927398 | 0.752885 |
|  |  |  | Enhancer_Hoffman.extend.500_0 | 0.153929 | 0.294083 | 0.331441 | 1.91051 | 2.153204 | 0.672381 |
|  |  |  | FetalDHS_Trynka_0 | 0.084757 | 0.586391 | 0.530831 | 6.918536 | 6.263011 | 0.343671 |
|  |  |  | FetalDHS_Trynka.extend.500_0 | 0.28501 | 0.410049 | 0.458539 | 1.438718 | 1.608851 | 0.78831 |
|  |  |  | H3K27ac_Hnisz_0 | 0.391169 | 0.409107 | 0.214996 | 1.045859 | 0.549623 | 0.933506 |
|  |  |  | H3K27ac_Hnisz.extend.500_0 | 0.422591 | 0.858798 | 0.299055 | 2.032221 | 0.70767 | 0.145846 |
|  |  |  | H3K27ac_PGC2_0 | 0.269477 | 0.713219 | 0.402386 | 2.646683 | 1.493212 | 0.273205 |
|  |  |  | H3K27ac_PGC2.extend.500_0 | 0.336034 | 0.480955 | 0.328905 | 1.431266 | 0.978784 | 0.65824 |
|  |  |  | H3K4me1_peaks_Trynka_0 | 0.171318 | 0.274478 | 0.577378 | 1.602152 | 3.370208 | 0.858145 |
|  |  |  | H3K4me1_Trynka_0 | 0.426568 | 0.780716 | 0.446849 | 1.830225 | 1.047543 | 0.439954 |
|  |  |  | H3K4me1_Trynka.extend.500_0 | 0.609157 | 0.604173 | 0.293772 | 0.991819 | 0.48226 | 0.986408 |
|  |  |  | H3K4me3_peaks_Trynka_0 | 0.041789 | 0.168317 | 0.349256 | 4.027753 | 8.357534 | 0.716096 |
|  |  |  | H3K4me3_Trynka_0 | 0.133307 | 0.716982 | 0.339529 | 5.378414 | 2.546962 | 0.076122 |
|  |  |  | H3K4me3_Trynka.extend.500_0 | 0.255482 | -0.29403 | 0.377819 | -1.15089 | 1.478847 | 0.116896 |
|  |  |  | H3K9ac_peaks_Trynka_0 | 0.03877 | -0.23207 | 0.354657 | -5.98576 | 9.147676 | 0.444881 |
|  |  |  | H3K9ac_Trynka_0 | 0.126111 | 0.789373 | 0.410177 | 6.259347 | 3.252506 | 0.089125 |
|  |  |  | H3K9ac_Trynka.extend.500_0 | 0.230583 | 0.442501 | 0.30732 | 1.91905 | 1.332791 | 0.504277 |
|  |  |  | Intron_UCSC_0 | 0.387453 | 0.555799 | 0.181554 | 1.434497 | 0.468583 | 0.331361 |
|  |  |  | Intron_UCSC.extend.500_0 | 0.39713 | 0.535948 | 0.142839 | 1.349552 | 0.359678 | 0.328584 |
|  |  |  | PromoterFlanking_Hoffman_0 | 0.008427 | -0.04378 | 0.176129 | -5.19461 | 20.89989 | 0.767216 |
|  |  |  | PromoterFlanking_Hoffman.extend.500_0 | 0.033471 | -0.11486 | 0.198178 | -3.43171 | 5.920801 | 0.444516 |
|  |  |  | Promoter_UCSC_0 | 0.031164 | 0.058569 | 0.209236 | 1.879386 | 6.714088 | 0.895386 |
|  |  |  | Promoter_UCSC.extend.500_0 | 0.038627 | 0.197321 | 0.156995 | 5.108351 | 4.064384 | 0.297335 |
|  |  |  | Repressed_Hoffman_0 | 0.461222 | 0.061012 | 0.520254 | 0.132283 | 1.12799 | 0.441205 |
|  |  |  | Repressed_Hoffman.extend.500_0 | 0.719053 | 0.394285 | 0.197363 | 0.548339 | 0.274477 | 0.091521 |
|  |  |  | SuperEnhancer_Hnisz_0 | 0.16842 | 0.521968 | 0.148543 | 3.099201 | 0.88198 | 0.018871 |
|  |  |  | SuperEnhancer_Hnisz.extend.500_0 | 0.171607 | 0.487953 | 0.136798 | 2.84344 | 0.79716 | 0.025348 |
|  |  |  | TFBS_ENCODE_0 | 0.132451 | 0.270263 | 0.588476 | 2.040483 | 4.442985 | 0.815614 |
|  |  |  | TFBS_ENCODE.extend.500_0 | 0.343444 | 0.519072 | 0.471425 | 1.511375 | 1.372643 | 0.711233 |
|  |  |  | Transcribed_Hoffman_0 | 0.345418 | 0.480785 | 0.431239 | 1.391892 | 1.248454 | 0.750559 |
|  |  |  | Transcribed_Hoffman.extend.500_0 | 0.763062 | 0.671393 | 0.269456 | 0.879868 | 0.353124 | 0.733177 |
|  |  |  | TSS_Hoffman_0 | 0.018219 | 0.013434 | 0.184548 | 0.737361 | 10.12949 | 0.979234 |
|  |  |  | TSS_Hoffman.extend.500_0 | 0.034826 | 0.093798 | 0.181524 | 2.693354 | 5.212392 | 0.742612 |
|  |  |  | UTR_3_UCSC_0 | 0.011054 | -0.01231 | 0.114901 | -1.11369 | 10.39429 | 0.838212 |
|  |  |  | UTR_3_UCSC.extend.500_0 | 0.026931 | 0.038184 | 0.127148 | 1.417851 | 4.721239 | 0.929034 |
|  |  |  | UTR_5_UCSC_0 | 0.005425 | -0.07372 | 0.113415 | -13.5894 | 20.90788 | 0.4659 |
|  |  |  | UTR_5_UCSC.extend.500_0 | 0.027806 | -0.27777 | 0.161663 | -9.98946 | 5.813983 | 0.02744 |
|  |  |  | WeakEnhancer_Hoffman_0 | 0.021093 | 0.02604 | 0.282699 | 1.234559 | 13.40279 | 0.985972 |
|  |  |  | WeakEnhancer_Hoffman.extend.500_0 | 0.088958 | 0.351986 | 0.293698 | 3.956747 | 3.30152 | 0.362349 |

**B: cell type-specific analyses**

| **BAG** | **Cell** | **Coefficient** | **Coefficient std error** | **P-value** |
| --- | --- | --- | --- | --- |
| GM | Oligodendrocyte | 1.31E-08 | 1.01E-08 | 0.096355 |
|  | Neuron | 7.36E-09 | 8.71E-09 | 0.199071 |
|  | Astrocyte | -1.1E-08 | 8.54E-09 | 0.901273 |
| WM | Oligodendrocyte | 2.47E-08 | 8.41E-09 | 0.001691 |
|  | Astrocyte | -1.6E-09 | 8.45E-09 | 0.57544 |
|  | Neuron | -9.2E-09 | 8.11E-09 | 0.872017 |
| FC | Astrocyte | 1.23E-08 | 6.59E-09 | 0.011238 |
|  | Neuron | 6.96E-09 | 6.31E-09 | 0.135152 |
|  | Oligodendrocyte | -4.9E-09 | 5.91E-09 | 0.796372 |

**eTable 6: Selected exposure variables for the forward Mendelian randomization**. We present here the traits, searching keywords, PubMed ID of the study, and the IEU ID. MR mimics randomized clinical trials using genetic variants (SNP) randomly allocated at conception as instrumental variables (IV) to estimate the causal effect of an exposure (e.g., alcohol consumption) on an outcome (e.g., GM-BAG). In essence, MR is less prone to confounding and reverse causation bias. Genetic variants, however, must be associated with the exposure variable (relevance assumption), not associated with the outcome biased by confounders (exchangeability assumption), and only associated with the outcome through the exposure (exclusivity assumption). In particular, we automatically queried these traits in the IEU GWAS database^7^ – curated GWAS summary statistics for MR – to extract the IVs from i) European ancestry, ii) non-UKBB studies (our GWAS were derived from UKBB data), iii) and large sample sizes. Another rationale for performing this hypothesis-driven MR analysis was the extensive coverage of UK Biobank (UKBB) in the IEU GWAS database, necessitating the exclusion of UKBB-based GWAS from our analysis to mitigate potential biases associated with sample overlap.

| Trait | Searching keyword | PubMed ID | IEU ID |
| --- | --- | --- | --- |
| AD | Alzheimer | 24162737 | ebi-a-GCST002245 |
| Breast cancer | cancer | 29059683 | ieu-a-1126 |
| Type 2 diabetes | diabetes | 22885922 | ieu-a-26 |
| Renin level | Renin | 33067605 | ebi-a-GCST90012038 |
| Triglyceride-to-lipid ratio | Triglyceride | 32114887 | met-d-XL_VLDL_TG_pct |
| AST | Aspartate aminotransferase | 29875488 | prot-a-1241 |
| BMI | Body mass index | 23563607 | ieu-a-85 |
